# Interplay of cis and trans mechanisms driving transcription factor binding and gene expression evolution

**DOI:** 10.1101/059873

**Authors:** Emily S Wong, Bianca M Schmitt, Anastasiya Kazachenka, David Thybert, Aisling Redmond, Frances Connor, Tim F. Rayner, Christine Feig, Anne C. Ferguson-Smith, John C Marioni, Duncan T Odom, Paul Flicek

**Affiliations:** European Molecular Biology Laboratory, European Bioinformatics Institute, Wellcome Genome Campus, Hinxton, Cambridge, CB10 1SD, UK.; University of Cambridge, Cancer Research UK - Cambridge Institute, Li Ka Shing Centre, Cambridge, CB2 0RE, UK.; University of Cambridge, Department of Genetics, Cambridge, CB2 3EH, UK.; Wellcome Trust Sanger Institute, Wellcome Genome Campus, Hinxton, Cambridge, CB10 1SA, UK.

**Author notes:** These authors contributed equally (Emily S Wong, Bianca M Schmitt). Co-corresponding authors: DTO, PF.

## Abstract

Noncoding regulatory variants play a central role in the genetics of human diseases and in evolution. Here we measure allele-specific transcription factor binding occupancy of three liver-specific transcription factors between crosses of two inbred mouse strains to elucidate the regulatory mechanisms underlying transcription factor binding variations in mammals. Our results highlight the pre-eminence of cis-acting variants on transcription factor occupancy divergence. Transcription factor binding differences linked to cis-acting variants generally exhibit additive inheritance, while those linked to trans-acting variants are most often dominantly inherited. Cis-acting variants lead to local coordination of transcription factor occupancies that decay with distance; distal coordination is also observed and may be modulated by long-range chromatin contacts. Our results reveal the regulatory mechanisms that interplay to drive transcription factor occupancy, chromatin state, and gene expression in complex mammalian cell states.

## INTRODUCTION

Understanding how genetic variation propagates into differences in complex traits and disease susceptibility is a major challenge. Evolutionary studies have revealed examples of regulatory variants linked to different organismal phenotypes^1^. Genome-wide studies have also found that many common disease-associated genetic variants lie in regulatory sequences^2–4^ with genetic changes at local-regulatory elements leading to coordinated chromatin changes within constrained genomic domains^5,6^.

A key determinant of transcriptional activation and spatiotemporal specificity is the affinity with which collections of transcription factors (TFs) bind to gene regulatory regions^7–9^. How TF binding specificity and strength is shaped by cis- and trans-acting variation remains poorly understood^10^, and understanding the interplay between TF binding and the surrounding chromatin state is critical for determining phenotypic diversity.

Cis-acting sequence changes substantially modulate TF occupancy^11,12^, but direct disruption of TF-DNA binding motifs is relatively rare^13–19^. This seemingly conflicting observation may be potentially explained by changes to surrounding chromatin state, long range TF-TF connectivity^6^ or cis-acting binding determinants near but outside the core binding motif^20^.

Strategies used to dissect cis- and trans-acting mechanisms include quantitative trait loci (QTL)-based analyses and F1 crosses of genetically inbred organisms. QTL analysis correlates a measured molecular trait, such as gene expression or TF binding intensity with genetic variation. However, fully distinguishing between regulatory divergence in cis and in trans in expression quantitative trait loci (eQTL) and chromatin immunoprecipitation quantitative trait loci (ChIP-QTL) studies^21^ requires large numbers of genetically diverse samples to achieve statistical power^22–25^. In addition, trans-effects are extremely difficult to identify and then validate^26,27^.

Alternatively, regulatory mechanisms can be revealed by analysis of the patterns of divergence occurring in F1 genetic hybrids; this approach has been widely used to analyze gene expression in yeast^28,29^, maize^30^, fruit flies^31–33^ and mouse^34,35^. By placing two alleles in a shared trans environment and comparing their allelic occupancy, the relative cis and trans contributions to a measured molecular trait can be evaluated^36^. Any variance from the occupancy observed in the parent F0 mice can be assigned to the influence of trans-acting variation. Like QTL-based approaches, analysis of F1 data results in a probabilistic description of the role of cis and trans effects. However, the use of F1 crosses classifies mechanisms underlying regulatory changes as either cis- or trans-acting more accurately than eQTL approaches, because the functional differences in vivo between the two alleles are directly evaluated in F1 mice^36^, rather than distance restricted searching between causative variants and TF occupancy differences as is common in eQTL approaches and which make distant eQTLs impossible to discover^37^. Formally, this system analyses the correlation between specific variants and observed functional effects, i.e. the effect of a variant on either cis- or trans-regulation. Although it is generally not possible to unambiguously assign causality from a specific variant to a functional effect, for simplicity in this study we will use the term ‘regulatory mechanism’ to refer to this connection.

Here we employ F1 hybrids to comprehensively dissect TF binding differences in mammals. We created first-generation genetic hybrids from divergent mouse sub-species to dissect trans-acting mechanisms that affect both chromosomes equally due to a shared nuclear environment, from the allele-specific differences caused by cis-directed mechanisms^32,33,38,39^. We leveraged this strategy to interrogate the inheritance of TF binding occupancy^40,41^. We find that changes to TF binding occupancy is predominately mediated by variation that acts in cis, and is thus additively inherited. In addition, cis-acting variation is able to influence multiple transcription factor binding sites (TFBS). Finally, we observe coordination in the regulatory mechanisms between TF binding occupancy, chromatin state and gene expression by incorporating matched transcriptomic data from RNA-seq^35^. Our results provide a comprehensive and quantitative overview of how different layers of regulatory variation create tissue-specific transcriptional regulation.

## RESULTS

### Transcription factor binding in mouse reciprocal crosses

In order to dissect the extent of cis and trans variation in TF occupancy variation, TFBS occupancy was mapped with six biological replicates using chromatin immunoprecipitation followed by sequencing (ChIP-seq) against three liver TFs (HNF4A, FOXA1, CEBPA) in inbred mouse sub-species *C57*BL/6J (BL6) and CAST/*EiJ* (CAST) and their F1 crosses (BL6xCAST and CASTxBL6) (**Figure 1a, Supplementary Figures 1–3, Supplementary Tables 1-2, Methods**); all data are in ArrayExpress (E-MTAB-4089). The large number (~19M) of single nucleotide variants (SNVs) between two parental strains, which are estimated to have less than 1 million years divergence^42^, is comparable to that found in human populations^43^, and permits a substantial proportion of allele-specific TF binding to be measured.

**Figure 1.**
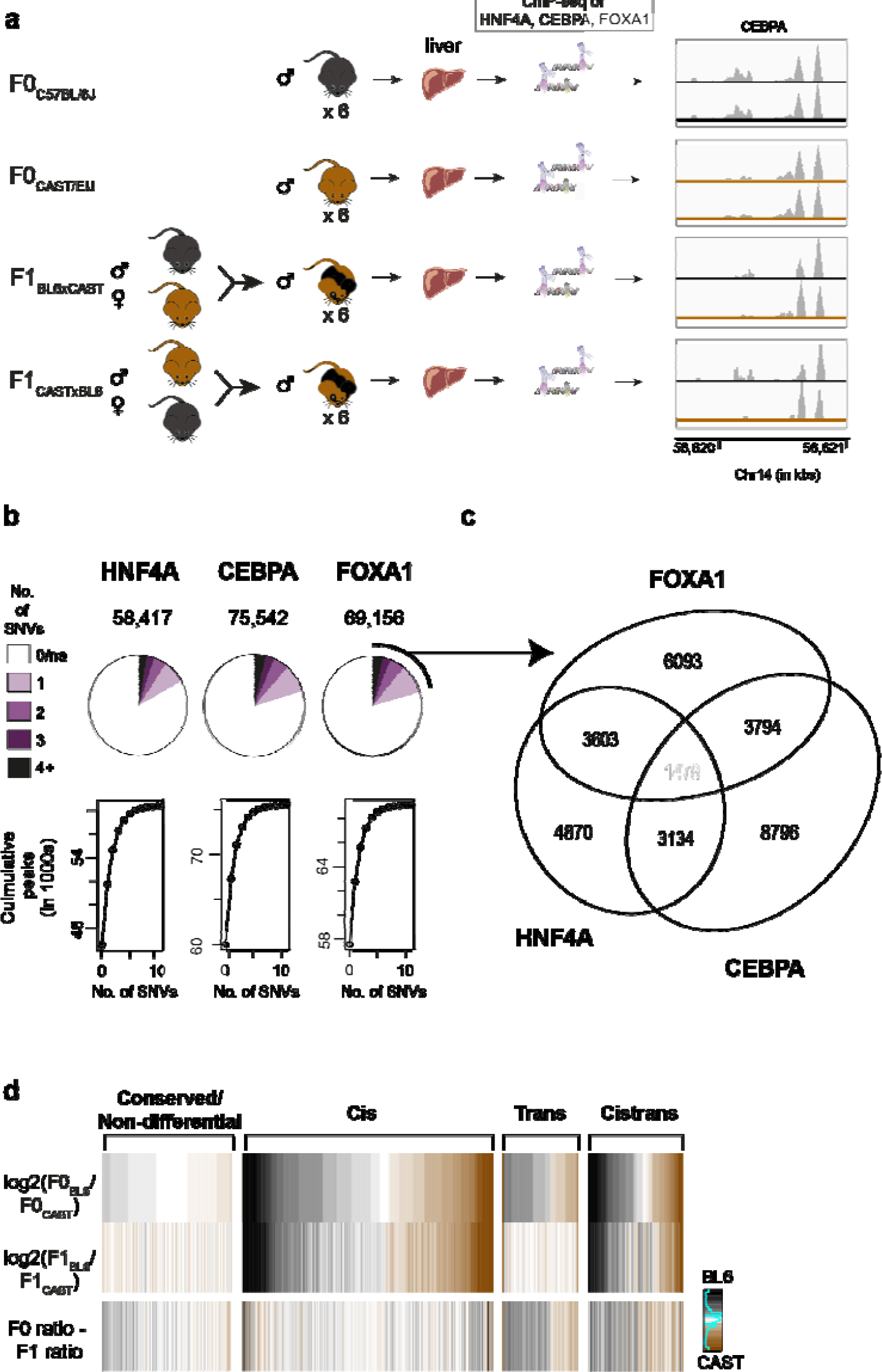
F1 mice were used to interrogate the regulation of TFBS variation. **(A)** *In vivo* binding of liver-specific TFs FOXA1, HNF4A and CEBPA were profiled in the livers of male mice from inbred strains C57BL/6J (BL6), CAST/EiJ (CAST) and their F1 crosses: C57BL/6J x CAST/EiJ (BL6xCAST) and CAST/EiJ x C57BL/6J (CASTxBL6). Six biological replicates were generated for each TF and genetic background combination. **(B)** The number of TFBS that could be classified with associated number of SNVs. **(C)** Venn diagram illustrates the numbers of classifiable SNVs that overlap between TFs. Each variant is at least 250bp from any other SNV. Numbers shown are the final numbers of regulatory loci used for downstream analyses. (**D)** Heatmap confirming overall accuracy of regulatory class assignments. BL6 (black) versus CAST (brown) binding intensity ratios for different regulatory categories for CEBPA. A subset of variants from each class was randomly sampled to match the overall distribution. Sparkline in key shows the number of observations at each color category where density is increasing from left to right.

Approximately 60-70,000 regions in the genome are bound by each TF (**Methods**), and approximately 20% had one or more SNVs with sufficient sequencing coverage to permit quantitative allelic resolution of TF binding (**Figure 1b**). Of these TFBS, in ~3-6% of these cases, SNVs directly disrupt a binding motif. Most (ca. 62%) SNVs are found in regions bound by only one TF, 34% are found in regions bound by exactly two TFs, and 5% by all three TFs, and are highly reproducible (**Figure 1c, Supplementary Figure 2**).

Cis and trans effects can be distinguished by the differences in binding affinities among F0 parents and their F1 offspring, as cis-acting variation must remain allele-specific^28,29,32,33,35^ (**Supplementary Figure 4a, Supplementary Figure 5**). TFBS that had informative SNVs for allelic resolution were classified into four regulatory categories – conserved (non-differential), cis, trans, and cistrans (affected by variants acting both in cis and in trans) (**Supplementary Figure 4b**) (**Methods**).

Differences in TF binding occupancies between the two mouse strains were most frequently affected by cis-acting variation (44-49%), followed by cistrans (14-17%) and trans (8-13%); 23-30% of TF binding was conserved despite the presence of one or more variants near the site of binding (**Supplementary Figure 4c**). Proportions of TFBSs belonging to each of the four categories were similar between all TFs. As expected, there are fewer conserved locations when SNVs directly disrupt the bound motif (**Supplementary Figure 4c**)^19^.

We confirmed the accuracy of the class assignment by visualising the difference in occupancy ratio between parental alleles and F1 alleles. By subtracting the F1 BL6:CAST ratio from the corresponding F0 ratio we found little difference in the allelic ratios from the parent and offspring in cis and conserved categories (**Figure 1d)**. In contrast, trans and cistrans categories show appreciable genotype specific signal. We validated our ChIP-seq measures of binding by performing allele-specific pyrosequencing (**Supplementary Figure 6**), confirming that approximately 40% of genetic variations that affect TFBS are cis-acting, compared with only 14% for liver-transcribed genes^35^.

### Characterization of TF binding occupancy

To quantitate the effect size of cis-acting variation on TF occupancy, we compared TF binding between F0 and F1 individuals using Pearson’s correlation (**Figure 2, Supplementary Figure 7**, **Methods**). In the absence of noise, a correlation coefficient of zero indicates that cis and trans contributions are equal, whereas a correlation coefficient of one indicates the absence of trans effects. We find Pearson’s r for TF binding to be significantly larger, and therefore cis dominated, compared to gene expression (TF binding: r=0.92, 95% CI (0.915, 0.919), P<2.2e-16; expression: r=0.62, 95% CI (0.607, 0.631), P<2.2e-16). Indeed, in 80% of instances when we compared any randomly chosen TFBS to any randomly chosen expressed gene, the magnitude of the cis effect was greater for TF occupancy than for gene expression (magnitude measured by the distance between F1 alleles over 10,000 random comparisons).

**Figure 2.**
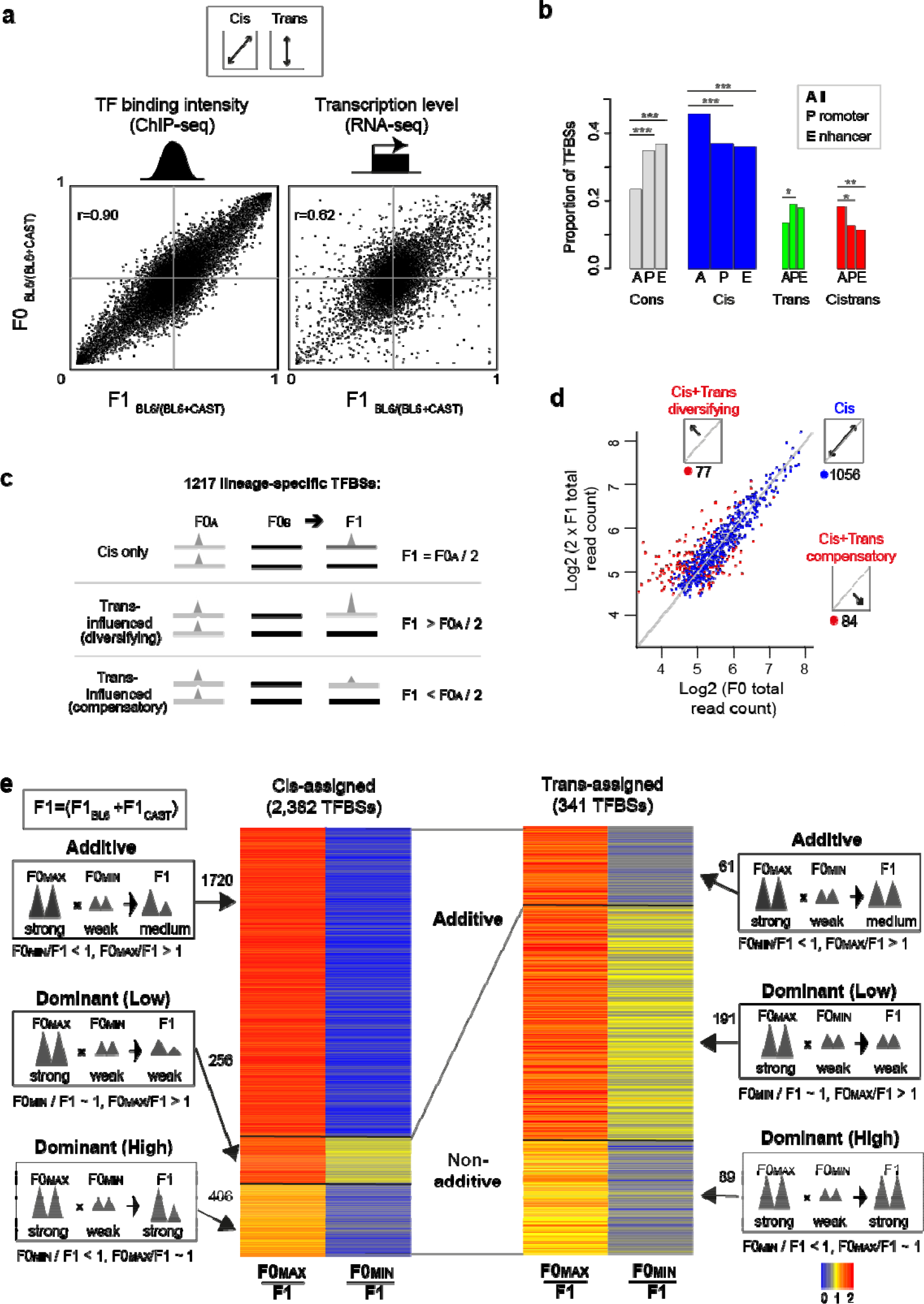
Differences in TF binding intensities strongly affected by variation acting in cis and are additively inherited. **(A)** Mean F0 versus F1 TF binding intensity ratios (BL6 versus CAST) for CEBPA are plotted in the left panel. The right panel shows mean F0 versus F1 gene expression ratios for liver-expressed protein-coding genes^35^. The correlation coefficient reflects the extent of cis-directed regulatory mechanisms. **(B)** Proportion of CEBPA binding locations at promoters and enhancers. The width of the bar is proportional to the overall number of TFBSs in the ‘All’ category. Binomial tests were used to test for enrichment at promoters and enhancers for each regulatory class based on the overall numbers of TFBSs (‘All’). ***P<0.0001 **P<0.001* P<0.05. **(C)** Most allele-specific TFBSs are affected by variation acting in cis. Lineage-specific TFBSs were defined as TFBSs where binding occurs either in BL6 or CAST in F0 individuals and in an allele-specific manner in F1 individuals based on a cut-off (F0_B6/(B6+CAST)_>0.95, F1_B6/(B6+CAST)_>0.95, F0_B6/(B6+CAST)_ < 0.05, F1_B6/(B6+CAST)_ < 0.05). These TFBSs can be sorted into the three categories described. (D) Mean CEBPA log2 F0 total read counts were plotted against mean log2 F1 read count (BL6 + CAST allele) multiplied by 2. For the scatterplot, we used averages across biological replicates. TFBSs affected by variation acting in cis are thus expected to fall along the diagonal and these have been colored blue (see C). Categories shown were determined by maximal likelihood estimation. (E) The majority of cis-directed TFBSs are inherited additively. TFBSs affected by variation acting in trans may show additive or dominant inheritance patterns in TF binding intensities. Different modes of inheritance were defined by comparing overall peak binding intensities between F0 and F1 individuals. Total F1 counts were individually scaled to 1 (yellow). Red indicates TFBSs where F1 > F0; blue indicates TFBSs where F1 < F0. CEBPA data is shown.

For lineage-specific TF binding locations, determined using *Mus spretus* as an outgroup^44^, we constructed statistical models to test the extent of variation acting in cis versus in cistrans to determine if variation that acts in cis or in trans is primarily responsibly for the creation of new TFBS. If the divergence was only due to variants acting in cis, the binding strength in the F1 allele will be half that in the F0 mouse. If TF binding in the F1 mouse were also influenced by variants in trans, then these binding intensities would be either greater or less than half the level found in the parent (**Methods**). The vast majority (87%, 1056/1217) of lineage-specific TFBS were affected by variation acting in cis (**Figure 2c-d**), while only 13% (161/1217) showed evidence of the influence of trans-acting variation. Overall, lineage-specific sites are up to two times less likely to have contributions from trans variants. Furthermore, we observed no lineage-specific TFBS were affected only by trans-acting variants (i.e. strain-specific in F0 but equally bound in F1). Our results strongly suggest that cis-directed variation either directly (e.g. modification of the binding motif) or indirectly (e.g. through remodelling of surrounding chromatin) play a required role in birth of TFBSs.

Next, we examined selective forces acting on TFBSs affected by variation in both cis and trans. Binding sites that show increased or decreased occupancy in the F1 due to cistrans-acting variation can have their effects decomposed into cis-acting variation that are either compensated by, or further changed by, trans-acting variation. We call changes compensatory when the difference in binding intensities in F1 < F0, whereas we call changes diversifying when the difference in F1 > F0. Under complete neutrality, both compensatory and diversifying trans effects should be equally favored^28^. Indeed, the frequency of compensatory versus diversifying effects is not significantly different at lineage-specific TFBSs (binomial test, P=0.6) (**Supplementary Figure 8a, Supplementary Table 3**), suggesting many allele-specific TF binding events are neutral. However, of the 2,563 CEPBA binding sites affected by variation in both cis and trans found on both alleles, 64% show compensatory changes (binomial test, P<2.2e-16), suggesting that shared TFBSs are more frequently subject to purifying or stabilizing selection (**Supplementary Figure 8a, Supplementary Table 3**). These numbers closely mirror the proportion of compensatory versus diversifying effects reported for gene expression in liver (68% compensatory, 32% diverging)^35^. No strain-specific TFBS affected only by variation in trans were observed (i.e. strain-specific in F0 but equally bound in F1). These results suggest that variation occurring is cis may either directly (e.g. by modification of the binding motif) or indirectly (e.g. through the opening up of chromatin by altering the shape of the DNA) play a required role in birth of TFBSs.

Additionally, we found little difference in selection pressure between strain-specific TFBSs that were gained and those that were lost in either BL6 or CAST lineages in the less than one million years^42^ since their divergence. Lineage-specific TFBS can be caused by: 1) lineage-specific loss of a TFBS that existed in the common ancestor of BL6 and CAST (plesiomorphic), or 2) lineage-specific gain since the most recent common ancestor in one strain (apomorphic) (**Supplementary Figure 8b**). To identify gained versus lost TFBS, we compared our lineage-specific TFBS with matched TFBS data obtained from livers of *Mus Spretus* (SPR)^19^, a mouse species of equal evolutionary distance (ca. ~1.5–2 MY) to both BL6 and CAST^44^. We distinguished between BL6 versus CAST lineage-specific binding sites that are apomorphic (present in BL6 not CAST or SPR and present in CAST not in BL6 or SPR) and plesiomorphic (shared between BL6 and SPR and between CAST and SPR but not BL6 and CAST). Around 35% of TFBSs strain-specific between BL6 and CAST were also found in SPR, placing a lower bound on the number of plesiomorphic TFBSs. Proportions of TF binding locations influenced by cis- or trans-acting variation were evenly distributed between apomorphic and plesiomorphic (binomial test, P>0.01) suggesting that there is little difference in selection pressure between strain-specific TFBS that are gained versus those that are lost.

We evaluated the potential regulatory activity of the TF binding by mapping the genome-wide locations of H3K4me3 (marking transcription initiation sites) and H3K27ac (marking potential enhancer activity)^45^ in F1 mouse livers (**Methods**). At promoters, TF occupancy changes affected by variation acting in cis and in both cis and trans were underrepresented (All TFs; binomial test; cis: P=1.1e-6, odds ratio (OR)=0.8; cistrans: P=1.2e-8, OR=0.6), and conserved sites were overrepresented (P<2.2e-16, OR=1.7) (**Figure 2b**). Regions showing enhancer activity were enriched for conserved TFBSs, and depleted for TFBSs that were directed by cis and cistrans variants (cis: P=3.4e-3, OR=0.8; cistrans: P=1.6e-6, OR=0.6; conserved: P=3.3e-8, OR=1.5).

The stability of genomic occupancy at TFBSs was assessed by evaluating the TF occupancy in BL6 mice with a single allele deletion of *Cepba* or *Hnf4a*, which can reveal regulatory activity and gene expression with more direct TF dependency^46^. When TF expression was reduced, the change in TF occupancy level was greater for binding sites influenced by cis-directed variation compared to those with a conserved binding pattern (**Supplementary Figure 9**). This suggests that TFBSs affected by variation acting in cis are more sensitive to changes in TF expression while non-differentially bound TFBSs are buffered.

TFBSs can be inherited in an additive or non-additive manner for variation that acts in cis or in trans^41^. Additive inheritance occurs when the combined occupancy of the F1 alleles is equal to the sum of the two parental (BL6 and CAST) F0 alleles^31,41,47^. Recall that cis and trans categories are defined by the occupancy ratio between parental alleles and F1 alleles (Methods), while inheritance concerns the total signal from both alleles. Dominant inheritance occurs when the total allelic occupancy in the F1 offspring is equal to that of either parent (**Figure 2e**). We fitted statistical models for both scenarios and evaluated them using Bayesian Information Criteria (BIC) (**Methods**).

Of the 2,382 TFBSs influenced by cis-acting variation (**Methods**), 72% (1,720) showed additive inheritance (of which 1,215 had BIC>2), whereas 28% (662) appeared dominant, which may partly reflect assignment errors (see **Discussion**). In contrast, of 341 TFBSs influenced by trans-acting variation 74% (280) exhibit dominant inheritance, whereas only 26% (61) were additive. Similar trends were observed for FOXA1 and HNF4A (**Supplementary Table 4)**.

We searched for evidence of over- and under- dominant patterns of occupancy inheritance that may correspond, respectively, to stronger or weaker F1 occupancy levels compared to parental measurements. In gene expression, this pattern of imbalance can be associated with hybrid incompatibilities^41^, and comprises approximately 27% and 8% (under- and over- dominant, respectively) of expressed genes between two strains of fruit flies^41^. In mice, we found that 6% and 11% of liver expressed genes showed under- and over- dominant modes of expression inheritance (**Supplementary Figure 10**). In contrast, less than 1% of sites in mouse tissues were determined as under- or over- dominant across all TFBSs (where BIC>1) (**Supplementary Table 5)**. Thus, under- or over- dominant TFBS inheritance appears rarely if at all.

In summary, variation in TF occupancy is strongly influenced by variation acting in cis, whereas TFBS affected by variation in trans are uncommon. In contrast to gene expression^41^, TFBSs are largely inherited additively, and TFBSs affected by variation acting in trans are mostly dominantly inherited (**Supplementary Figure 10**).

### Influence of cis-acting variation rapidly decays with distance

We asked what impact cis-acting variation have on TF occupancy at varying genomic distances because chromatin state can depend on distal functional elements^5,6^. For example, in humans eQTLs are considered local if they are within 2 Mb of the gene they influence^48^ and many distant eQTLs are known to exist^37^.

We first confirmed that overlapping binding events from different TFs share cis-acting variation more often than expected by chance (**Supplementary Figure 11**). We quantitated the affinity with which cis-acting variation influence distant TF binding occupancies using a complementary strategy to Waszak et al.^6^. Although the exact location of each causal variant is unknown, the genomic span (or effect distance) of a cis-acting variant can be inferred by examining co-variation in binding occupancies between neighbouring TFBSs (**Methods**, **Figure 3a, Supplementary Figure 12**).

**Figure 3.**
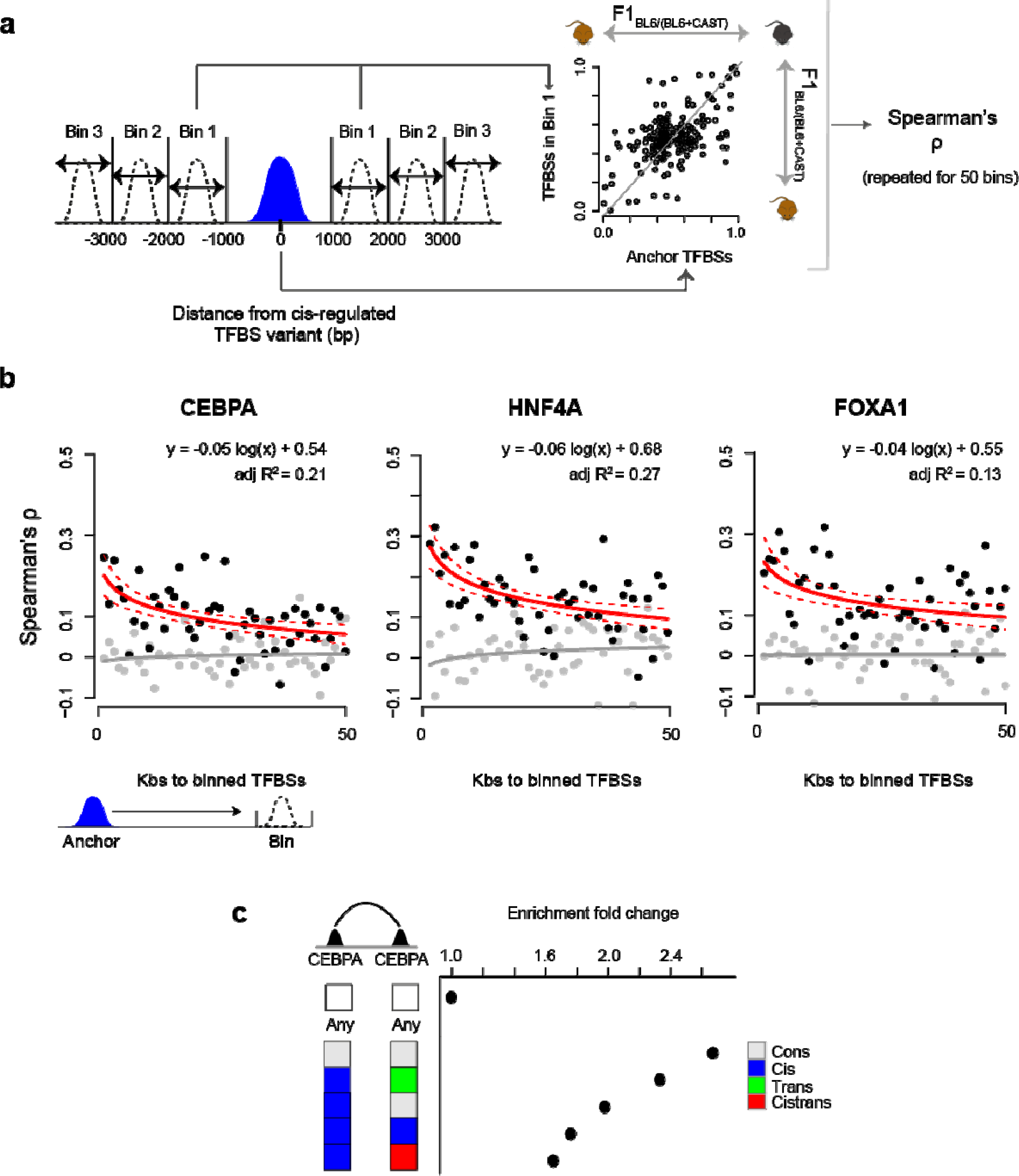
Effect of genomic distance on cis-acting inter-peak correspondence. **(A)** Strategy for measuring the span of cis regulatory effects. Successive 1kb bins were taken from each TFBS affected by variation acting in cis starting 400bps from the location of the SNV and extending in both directions. For each bin, Spearman’s ρ was calculated using the BL6:CAST allelic ratio between queried TFBSs against TFBSs assigned as anchorages for the analysis. **(B)** Spearman’s ρ values for each bin were plotted for each TF. The linear regression line (solid red) calculated from these values is shown. Red dashed lines mark the 90% confidence intervals of the true slope of the line. Grey dots represent the null background distribution of correlation values constructed by the random subsampling of TFBSs to anchor TFBSs (see **Methods**). The numbers of TFBSs in each randomly sampled bin were matched to those in the observed bins. The grey line is the linear regression line for the correlation values derived from sampled points. **(C)** TFBSs are enriched at regions of chromatin contact. Enrichment values were calculated compared with expected rate of chromatin contact given the general enrichment for contact in each regulatory dataset (i.e. cons, trans, cis, cistrans). ‘Any’ denotes the null background set of randomly chosen locations in the genome

The correspondence between TF binding occupancies decreases at a logarithmic rate, with similar trends observed across all three TFs (**Figure 3b)**. For example, the correspondence is 2-3 times lower at 50kb than at 3kb, but we nevertheless detected correspondence affected by variation acting in cis slightly above genomic background levels up to 400kb away (Rho=0.01–0.02, linear regression). We estimated using vector projection that the observed correspondence between TFBSs falls off relatively quickly for approximately 13kb and more slowly thereafter (**Methods**) suggesting that the genomic scope of a cis-acting variant on TF binding is on the order of 10kb. Our results were consistent across several bin sizes grouping nearby SNVs (**Supplementary Table 6**). Different TF binding locations appear to be similarly correlated, as shown recently for chromatin domains^5,6^.

Long-range coordination of TF occupancy could be affected by cis-variation via three-dimensional interactions, and we therefore searched for direct evidence that spatially distinct TFBSs interact (**Figure 3c**). We analyzed Hi-C data from BL6 mice^49^ to identify the interaction endpoints that overlap CEBPA binding locations (**Methods**). As expected, conserved sites were more likely to overlap long-range interaction endpoints (logistic regression: P<0.05, OR=1.14–1.20) (**Supplementary Table 7**). Chromatin interactions anchored on a cis-associated location were strongly enriched over the any-versus-any background (binomial test; P-value: cons versus cons=2.0e-8, cis versus trans=1.8e-9, cis versus cons=4.0e-10, cis versus cis=5.7e-6, cis versus cistrans=4.5e-4). Significant enrichment over the any-versus-any background set was observed for all categories of TFBS.

Our data support a model where the cis-acting variants causal for differences in TF binding occupancy are mostly proximal to the TFBS they affect. However, regions with TF occupancy, including TFBSs affected by variation that acts in cis, are disproportionately found at interaction endpoints for genomic domains, providing a possible mechanism for the observed long-range correlations.

### Coordination of regulatory mechanisms

The connection between genetic variation with TF binding, chromatin state and gene expression has recently been studied in human cell lines^14,15,17^. However, how genetic variants affect the interplay and temporal ordering of these regulatory layers remains poorly understood.

As above for transcription factor binding, we classified the regulatory mechanisms of variation underlying the allelic changes in chromatin state and transcription based on whether these differences are influenced by variation acting in cis, conserved, influenced by variation acting in trans, or by variation acting in both cis and trans (**Figure 4, Methods**). We then used logistic regression to establish whether the mechanisms responsible for regulating TF binding differences are enriched or depleted within the corresponding chromatin and gene expression categories.

**Figure 4.**
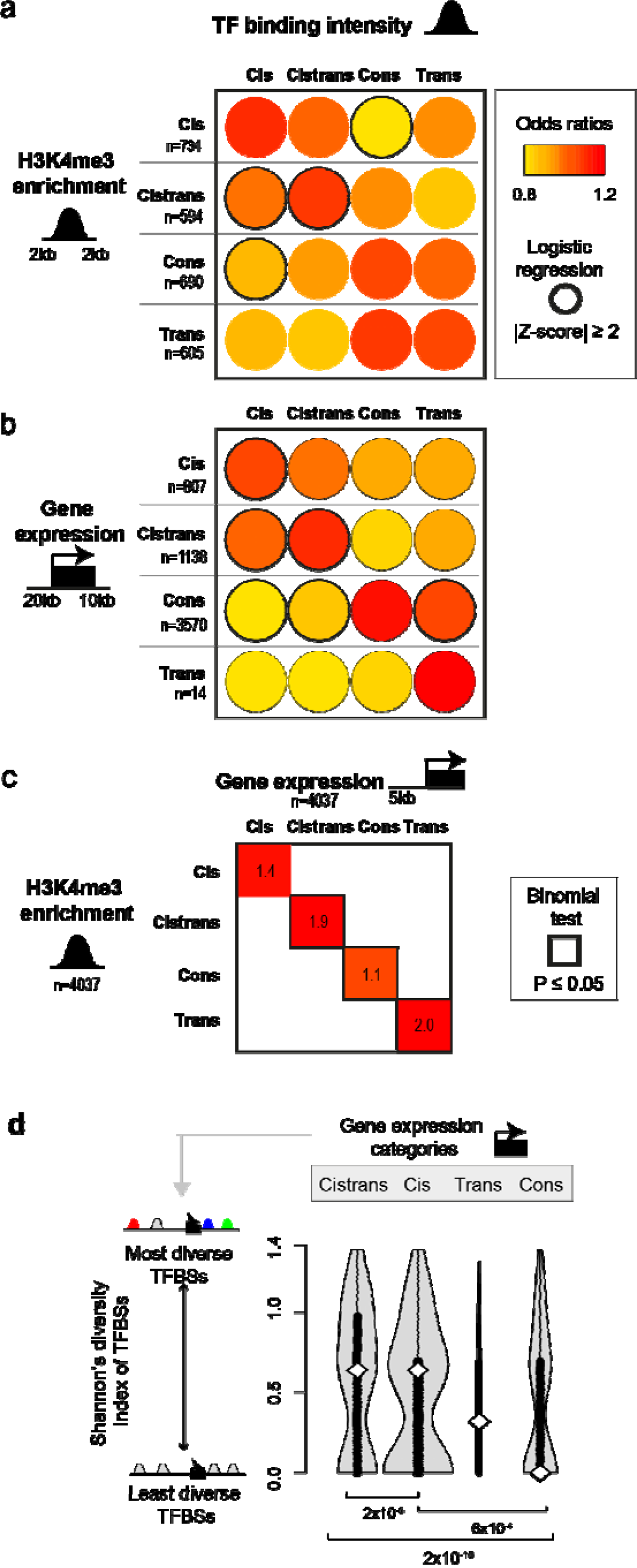
Genetic and epigenetic influences that change TF binding have parallel consequences for gene expression and chromatin. **(A-B)** Coordination between the regulatory categories of variation in TF binding occupancy variation and chromatin (A) and gene expression (B). Locations of the considered TFBSs are noted in the cartoons on the left. Separate logistic regressions were performed for each chromatin regulatory class (see Methods). Odds ratios were mean-centred for comparison across chromatin regulatory classes. Absolute values of Z-scores greater than two (α<0.05) are denoted by a black border. **©** Direct association between chromatin and gene expression. Genes were linked to H3K4me3 modifications if the mark was located within 5kb upstream of the TSS. Binomial tests were performed based on the expected background probability of observing the same regulatory mechanism underlying both expression and histone enrichment change. **(D)** High diversity in regulatory mechanisms of TF binding variation is associated with gene expression influenced by cistrans-acting variation. Calculations are on a gene-by-gene basis for TFBSs 20kb upstream and 10kb downstream of TSSs. These scores were compared between genes grouped by transcriptional regulatory class. Significant P-values for Mann-Whitney U tests are shown. The surface area of the violin plot is proportional to the number of genes in each class.

We found similar variant classes underlying TF binding occupancy, chromatin state, and gene expression at the same locus as represented by the darker red circles on the diagonal in **Figure 4a-b**. For example, the genetic variants that influence TF binding in a cis-acting manner are more likely to be collocated with H3K4me3 regions also showing changes influenced by cis-acting variation (top left red circle in **Figure 4a**). This is compatible with models proposed by Kilpinen et al.^15^.

Furthermore, there is a positive, albeit modest, correlation between the *direction of effect* between allelic changes in TFBS occupancy and gene expression (**Supplementary Figure 13**). In other words, when a TFBS increases its occupancy, then nearby gene transcription often increases (binomial test, P=2.9e-4) and with similar magnitude (Spearman’s rank correlation, rho=0.29, P=6.3e-12).

We controlled for the possibility that these effects are caused by differences in expression levels by using an alternative strategy that subsampled genes with least one TF binding event in the region 20kb upstream and 10 kb downstream of the TSS so that they had matched expression levels between regulatory classes. We found that genes showing conserved expression levels were depleted for TFBSs with occupancy affected by variation in cis (Mann-Whitney U test on a per gene basis comparing the numbers of different TFBSs near conserved regulated genes against genes where expression is influenced by variation in cis or cistrans, P=9.8e-11 and 2.2e-16, respectively). Hence, genes whose expression variation is influenced by variation in both cis and cistrans possessed a higher than expected number of TFBSs proximal to the TSS that were influenced by variation in cis. Analysis of genes with expression affected by trans variants was uninformative due to the small number of genes (14) in this category.

We also confirmed previously observed correlations between promoter chromatin state and gene expression^48^. We identified a subtle but significant correspondence between the types of regulatory variation underlying promoter activity differences and gene expression differences (binomial test; cis P=0.04, cistrans P=0.03, conserved P=0.02, trans P=0.25) (**Figure 4c**). These results suggest robustness to our overall analysis.

Finally, TFBSs often act in concert with one another. Using Shannon’s entropy, we compared the mechanistic diversity of TF binding variants with the regulatory mechanisms of variation affecting nearby gene expression (**Methods**). In effect, this analysis asks whether the collective effect of the cis- and trans-acting variation underlying changes to occupancy levels of multiple TFBSs can propagate to gene expression. Expression influenced by variation in cis or both cis and trans was significantly more likely to be associated proximally to TFBSs that are themselves affected by variation acting through diverse mechanisms (Mann Whitney U test) (**Figure 4d**). In contrast, conserved expression was likely to be associated with TFBSs directed by a similar type of variant. We controlled for the possibility that the association between the diversity of TFBS and the category of gene expression might be due to differences in gene expression levels within each category by repeating the analysis using expression matched subsets of genes from each regulatory category. We observed little difference on our core results (Mann Whitney U test; cistrans versus conserved: P=1.4e-7, cis versus conserved: P=1.4e-3). Our results therefore suggest that genes affected by variation that is cis-acting and cistrans-acting are more likely to be proximal to TFBS of high mechanistic diversity.

## DISCUSSION

Directly connecting genome-wide observations of transcription factor binding with functional outputs in gene expression is challenging because of what appears to be two conflicting observations. On the one hand, most variation in the human genome associated with complex disease and other phenotypes is non-coding^4^. Even for Mendelian disorders, exome sequencing can suggest causative sequence changes in only a minority of cases (~25%)^51^. Both point to a major role for functional sequence changes in the regulatory regions of the genome, which subsequently lead to changes in gene expression. On the other hand, TF binding demonstrates both variability between even genetically identical individuals and such strikingly rapid evolutionary change^52^ that it is tempting to conclude that the vast majority of TF binding is non-functional “biological noise”^53^.

Here, we have undertaken a detailed and comprehensive dissection of the genetic mechanisms driving TF binding occupancy differences in mammals and integrated these results with chromatin and gene expression information. Our initial findings regarding how genetic sequence variation associates with TF binding differences between alleles are consistent with previous reports at a more limited set of locations in murine immune cells^54^, human lymphoblast cells^14,18^, and using computational simulations^55,56^. Specifically, almost three-quarters of assayed quantitative differences in TF binding occupancy appear to be the result of nearby genetic differences that acts in cis.

However, our integrated analysis extending from TF binding to output gene expression using F1 inter-strain mouse crosses revealed a number of novel insights. First, the vast majority of trans-directed TF binding differences are dominantly inherited. Although most binding influenced by cis-acting variation is inherited additively, as expected, a small proportion appears to show dominance/recessive inheritance. One plausible biological explanation is the presence of variation acting in trans that does not interact with cis-acting variation at each allele. Despite this, cis and trans-acting variation driving TF occupancy change show clear differences in their mode of inheritance. Second, allelic differences in TF binding are correlated at kilobase distances above the genomic background, likely influenced by neighbouring cis-acting variation. A minor fraction of TFBSs show long-range coordination, which may be driven by enrichment of TFBS at chromatin contacts. Such long-range correspondence is similar to recently described coordination of chromatin states within topological domains^6,57^. Third, we demonstrate interplay between the different mechanisms of variation that underlie transcription factor binding and tissue-specific gene expression *in vivo*. Aspects of the regulatory interplay between chromatin and gene expression has been reported in human cell lines and mouse species^15,58–61^.

The F1 genetic cross analysis is very effective at disambiguating cis- and trans-acting regulation overall. Our data shows that genetic variants can (simultaneously) direct TF binding, chromatin, and gene expression changes using a similar combination of regulatory variation that acts in cis and trans. However, the full temporal order of regulatory events cannot be determined from our data. For instance, our results do not reveal whether genetic variants first affect TF binding which then affects chromatin - or vice versa. However, the presence of an additional trans component in gene expression suggests that it is downstream of both TF binding and chromatin modifications.

The independently determined categories of regulatory variation correspond well between TF occupancy and gene expression. This is potentially surprising given the difference in the overall regulatory repertoire between TF binding and gene expression. Namely, protein-DNA interactions are shaped by a comparatively simple combination of DNA sequences, chromatin context, and (in some cases) noncoding RNA associations. In contrast, a multitude of regulatory processes influence gene expression, including TF binding as well as post-transcription processing, translation rate and mRNA degradation. Our results support a model whereby the variation underlying gene expression differences arise substantially from a composite of the variation that modulate TF binding differences in multiple individual TFBSs.

Our analysis has specific limitations. Our approach cannot analyse the majority of TFBSs where no informative SNV is present, and these unclassified TFBSs are more likely to be conserved. However, a change in the relative proportion of regulatory categories is not expected to influence our key findings, which were focused on the regulatory mechanism effect size. Our analysis ignores structural variants, and we have not directly measured fitness in the F1 animals. We also cannot preclude the possibility that tissues other than liver may demonstrate a greater affect of trans-acting variation on TF binding differences. Although most tissue-specific gene expression appears to be driven by combinatorial TF binding of dozens of TFs^10^, we have profiled only a subset of three. However, analysis of the occupancy of over a hundred TFs in one tissue strongly suggest that our data will reflect the typical mechanistic contributions influencing the evolution of all tissue-specific TFs^62^. Finally, our technical definition of the binding sites affected by both cis and trans variation will include TFBSs with high biological and/or technical heterogeneity.

Our work builds upon previous findings of genomic coordination among TF binding, chromatin marks and transcription^5,6,15,63^ and highlights the key role played by the basal variation that underlie TF binding in directing regulatory change.

## METHODS

### Sample collection and preparation

All mice were housed in the same husbandry conditions within the Biological Resources Unit in the Cancer Research UK-Cambridge Institute under a Home Office Licence. C57BL/6J (stock Number 000664, imported from Charles River Labs) and CAST/EiJ (stock number 000928, imported from The Jackson Laboratory (www.jax.com)) mouse strains were used in experiments as parental strains (F0) as well as for breeding of reciprocal crosses of F1 mice. All mice used in the experiments were males between eight and 12 weeks of age, and harvested at the same time of day (between 8 and 11am). Liver perfusion was performed on mice post mortem, prior to tissue dissection. Harvested tissues were formaldehyde cross-linked for ChIP-seq experiments. Before cross-linking, dissected tissue was immediately chopped post mortem and added to a cross-linking solution containing 1% formaldehyde. Tissue was incubated for 20 min prior to quenching with 1/20th volume of 2.5 M glycine. Samples were incubated for a further 10 min before washing with PBS and flash-freezing and storage at —80°C.

### Generation of HNF4A and CEBPA heterozygous mice

To create HNF4A and CEBPA heterozygous knockout mice, we acquired mice with targeted alleles from The Jackson Laboratory (HNF4A stock number: 004665^64^; CEBPA stock number: 006230^65^). Heterozygous knockouts were generated via the Cre-loxP system^66^ using the germline deleter strain PgkCre^67^, obtained from The Jackson Laboratory, and crossing it to *Cebpa^FLOX/FLOX^* and *Hnf4a^FLOX/WT^* mice. Ear biopsies were taken at the time of weaning for genotyping to confirm deletion via PCR (**Supplementary Table 8**).

### ChIP-seq experimental procedure

The ChIP-seq protocol used was as described by Schmidt et al.^68^. Briefly, livers were isolated from 10 to 12 weeks old mice and liver tissue was post-mortem cross-linked using 1% formaldehyde (v/v), lysed and sonicated. Protein-bound DNA was immunoprecipitated using 10µg of an antibody against CEBPA (Santa Cruz, sc-9314), HNF4A (ARP 31946_P050), FOXA1 (ab5089, Abcam), H3K27ac (ab4729, Abcam), or H3K4me3 (Millipore 05-1339). Immunoprecipitated DNA was end-repaired at 20°C for 30 min, Adenine overhang was added at 37°C for 30 min, and Illumina sequencing adapters ligated at room temperature for 15 min before 16 cycles of PCR amplification. PCR conditions: 1) 98°C – 30 sec; 2) 98°C – 30 sec, 65°C – 30 sec, 72°C – 30sec, 16 cycles; 3) 72°C – 5 min. DNA fragments ranging from 200- to 300-bp in size were selected on a 2% agarose gel for 50-bp single-end read sequencing on an Illumina HiSeq 2000 according to the manufacturer’s instructions.

### Validation of allele-specific TF binding with pyrosequencing

We performed pyrosequencing to confirm the allele-specific occupancy of CEBPA in livers from F1 mice in both genetic cross directions. The assays and primers (**Supplementary Table 9**) for pyrosequencing were designed using PyroMark Assay Design Software. The annealing temperature for PCR primers was optimized by gradient PCR. Primers’ efficiency was confirmed using quality controls with different proportion of BL6 and CAST DNA (0/100%, 30/70%, 50/50%, 70/30%, 100/0%). PCR conditions: 1) 95°C – 5 min; 2) 94°C – 30 sec, optimized t°C – 30 sec, 72°C – 55sec, 40 cycles; 3) 72°C – 5 min. PCR product was mixed with streptavidin beads dissolved in binding buffer and gently shaken for 20 min. Sequencing primers were dissolved in annealing buffer and aliquoted into PSQ plate. DNA-Beads were cleaned on the PyroMark vacuum workstation and then mixed with PSQ Primer/Annealing Buffer. The samples were incubated at 85°C for 3 min, centrifuged for 3-4 minutes at 2500 rpm and then loaded to the pyrosequencer. PyroMark Gold Q96 SQA Reagents were used to load the pyrosequencer.

### Estimation of allele-specific binding level

We constructed the *Mus musculus castaneus* genome assembly using CAST/EiJ SNV calls (ENA accession: ERS076381) against the *Mus musculus* reference assembly (C57BL/6J)^69^. Single nucleotide variants (SNVs) were mapped from their original calls on NCBI37/mm9 to the latest version of the mouse assembly, GRCm38.p2/mm10, and nucleotides at each base position were changed to reflect point mutations in CAST. SNV calls were available for all autosomes and the X chromosome.

To assess allele-specific binding and histone enrichment, we aligned reads to an alignment index comprising of both GRCm38.p2/mm10 (BL6) and CAST assemblies. Indexing of the genomes was performed using BWA (Version 0.7.3a)^70^. Raw sequencing reads were first filtered and trimmed using Trimmomatic (Version 0.3)^71^. We required a minimum phred score of 30 using a sliding window of 20 bps, and only kept a read if it matched these criteria while maintaining a minimal overall length of 40bp. We aligned filtered reads using BWA with a maximum of 2 mismatches per read (-n 2). Reads that mapped equally well to multiple locations were discarded by filtering based on the ‘XT:A:U’ alignment tag. Our alignment statistics showed our approach assigned reads to each strain with high specificity (see **Supplementary Figure 3**). The proportion of F1 reads aligning to the combined BL6 and CAST genomes was roughly 51:49, respectively. Proportions of BL6 TFBSs versus CAST TFBSs called from these alignments were similar.

The mpileup program from the SAMtools package^72^ was used to count the number of reads that overlapped each base of the joint assembly. We then filtered these counts to retain only those genomic locations where it was possible to distinguish between BL6 and CAST backgrounds. We only retained sites for analysis where a minimum of 10 reads mapped to either F0 CAST or F0 BL6 across replicates. For F1 crosses, we retained sites overlapping at least 10 reads for at least 10 allele-specific replicates. We repeated these steps on a site-specific manner for each TF/histone mark, irrespective of whether multiple SNVs existed at each ChIP-seq peak.

Prior to fitting statistical models and further downstream analyses, we normalized for sequencing depth by adjusting for differences in library sizes across biological replicates in F0 and F1 populations for each TF/histone mark. A constant scaling factor was estimated for each library based on the median of the ratio of reads at each SNV over its geometric mean across all libraries tested. This normalization constant was then applied to each library under the assumption that count differences attributable to biological effects only exists in a small proportion of the total number of sites. This procedure was performed using R Bioconductor package ‘DESeq’^73^.

To assess overall peak counts and determine the quality of each ChIP experiment, we also aligned reads from each library (F0 and F1) to the GRCm38.p2/mm10 genome using GSNAP^74^ with a less stringent mapping criteria. We used a less conservative mismatch threshold (maximum mismatch of 3 bases per read) to allow F1 reads derived from the CAST allele to map against the BL6 genome. Based on overall SNV numbers between the strains, a rough estimation suggests that there are approximately 1 SNV every 100 bps, which distinguishes the strains. Regions bound by both TFs and covalently modified histones were called using MACS1.4^75^ using default parameters.

To mitigate the impact of potential batch effects, biological replicates for each TF for each genetic background were prepared and sequenced in three independent flowcells.

We estimated TF occupancy levels for the histone modification H3K4me3 by taking into account the fact that histone marks typically localize over a broader genomic region than do TFBSs. Wider regions cause a dilution in the number of reads overlapping SNVs, relative to binding site numbers and sequencing depth. Hence, to increase our ability to resolve binding differences at H3K4me3 loci, we summed the counts of all SNVs overlapping the same region. To ensure background-specific peaks were captured, we constructed a summary peak file comprised of the union of genomic intervals from peak calls from individuals of different genetic backgrounds (BL6, CAST and BL6xCAST) (library reference: do3342, do3337, do3411).

We identified between 6,000-8,000 TF bound regions per TF where two or more SNVs lie within close (<250 bp) proximity; ~85% of these co-located SNVs showed the same allelic direction of TF binding between BL6 and CAST. To avoid multiple counting of TF binding events, we only used one SNV in any 250 bp region in further analyses. Our results were highly reproducible among replicates (**Supplementary Figure 2**) with similar numbers of reads mapping to each genome (**Supplementary Figure 3**).

### Statistical models for identifying regulatory mechanisms

ChIP-seq read counts were used as a proxy for the binding intensities of a TF to the DNA^7^. Sites were classified into regulatory categories using the method of Goncalves et al.^35^.

We defined as <u>conserved</u> those regions with equal TF binding occupancy between BL6 and CAST in both F0 and F1 individuals, despite the presence of one or more variants near the site of binding; these types of sites could also be described as non-differentially bound^28^. We defined TFBSs influenced by cis-acting variation as sites where the TF occupancy ratios between BL6 and CAST genomes found in the F0 parents is the same as that observed between alleles in the F1 offspring, meaning that binding occupancy differences between strains were determined by locally acting genetic sequences. We defined TF binding influenced by trans-acting variation based on TF binding occupancy differences between parents, but not between alleles in the F1 offspring. Finally, we defined binding sites influenced by cistrans-acting variation as showing a complex mixture of cis and trans acting variation.

For each TF or histone mark, F0 counts from each strain were modelled as a negative binomial marginal distribution, while F1 counts were modelled using a beta-binomial distribution where the parameters of the beta distribution modelled the proportional contribution from each allele. For each TF and histone mark, there were 6 replicates (*i*) for each F0 strain and 12 replicates (*j*) for F1 samples. F0 counts for each strain (*x_i_*, and *y_i_*) were assumed to follow negative binomial distributions while F1 counts (*n_j_*), were modeled on an allele-specific basis (*z_j_*) using a beta-binomial distribution:

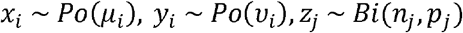

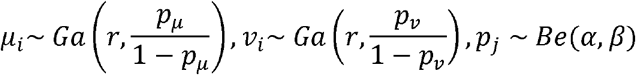

where *x_i_* is formally defined as the binding intensity of the variant in the *i*th C57BL/6J F0 mouse, *y_i_* is the binding intensity of the variant in the *i*th CAST/EiJ F0 mouse, *n_j_* is the number of reads mapping across both allelic variants in the *j*th F1 hybrid and *z_j_* is the number of reads mapping to the C57BL/6J allele in the *j*th F1 hybrid.

We estimate the dispersion parameter *r* for F0 samples using the ‘estimateDispersions’ function within ‘DESeq’ with local regression fit. *r* was used as the reciprocal of the fitted dispersion value from ‘DESeq’.

We constrained parameter estimation for each distribution based on four different regulatory scenarios and derived maximum likelihood values for each hypothetical case on a site-by-site basis. The four models are described below:

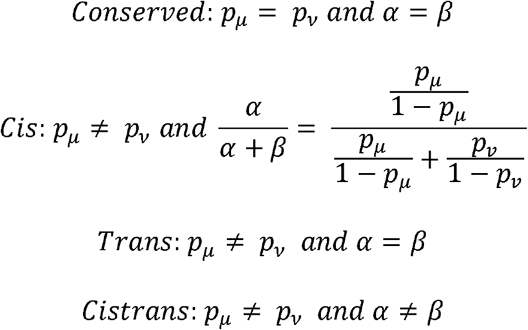

To identify the most probable model at each variant we used the Bayesian information Criterion (BIC).

To avoid confounding results from the analysis of variants derived from the same binding site, downstream analyses only used variants spaced at least 250 bps apart. Hence, where two or more variants were found spaced within 250bps of one another, only one variant was chosen for subsequent analyses.

### Identification of motif-disrupting variants

MEME^76^ was used to perform *de novo* search for enriched motifs for each TF using one randomly chosen ChIP-seq library per TF (library identifiers do3488, do3463 and do3483). Sequences +/-50bp from all peak summits were extracted for analysis; where multiple motifs exist in a peak, the motif sequence with the best score was retained.

### Regional enrichment of mechanisms driving TF occupancy

Enrichment for TF regulatory categories that overlapped the location of histone marks was assessed using the exact binomial test. Colocation was defined using an overlap of 1bp. The probability of success in the Bernoulli trial was defined for each TF based on its proportion of binding categories.

To assess whether co-locating TFs (i.e. binding at the same SNV) share the same regulatory category (i.e. cis, cistrans, conserved, trans) more often than expected by chance, we calculated the expected probability of Bernoulli success as follows:

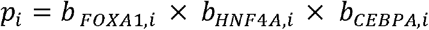

where *b* is the proportion of TFBSs in regulatory category *i* at TFBSs where all three TFs co-locate.

### Differential binding analysis of heterozygous versus WT mice

The genome-wide binding of CEBPA, HNF4A and was assessed in *Cepba^FLOX/-^* and *Hnf4a^FLOX/-^* mice. Three biological replicates per condition (HET or WT) per antibody were compared to quantify changes in TF binding intensity after heterozygous TF deletion. We then sorted the TFBSs based on whether their occupancy was conserved, or affected by variation in cis or both cis and trans. Binding intensities were considered as the number of reads at the summit of peaks that were called by MACS1.4^75^. The same WT input libraries were used for peak calling in both HET and WT samples. We filtered out peaks with a read count cut-off of less than 11 reads in less than 5 libraries. Prior to differential binding comparisons, upper quantile normalization^77^ was used to adjust for differences in sequencing depth between libraries. For each TF, ‘edgeR’^78^ was used to identify peaks with different binding intensities between HET and WT samples, using a significance cut-off of FDR<0.1.

### Assigning modes of TF occupancy inheritance

To identify the mode of inheritance of TF binding intensities at non-conserved TFBSs, F0 and F1 libraries were first adjusted for differences in sequencing depth using the median of the ratio of reads at each SNV over its geometric mean across all libraries as a constant normalization factor for each library^73^. Next, data from each SNV was fitted to statistical models reflecting either additive or dominant/recessive inheritance patterns. Models were constructed based the following premise: if offspring binding intensities were inherited via an additive mode of inheritance, we would expect the combined offspring binding intensity from both alleles to equal the summed binding intensity of parental alleles; on the other hand, if inherited through a dominant/recessive mode of inheritance, we would expect the combined binding intensity in the offspring across both alleles to equal the total intensity of one but not the other of its parents. We assumed read counts followed negative binomial distributions. Here, we formally define the models:

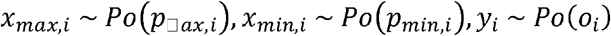

*x_max,i_* is defined as the normalized read count binding intensity of the variant in the *i*th F0 mouse from the parental strain showing the higher median binding intensity among replicates, *x_min,i_* is the normalized read count binding intensity of the variant in the *i*th F0 mouse from the parental strain with the lower median binding intensity among replicates. *y_i_* is the binding intensity of the variant in the *i*th F1 mouse summed across both alleles.

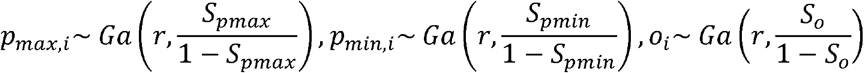

As above, the dispersion parameter, *r*, was estimated using ‘DESeq’. We used maximum likelihood estimation to fit the counts to the models below and used BIC to assess which of the following two models best fit counts from each site affected by variation in cis or trans.

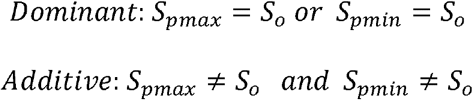

We excluded those sites from our results where the parameter estimated for the offspring, *S_o_*, was indistinguishable from the parameters estimated for both parent, i.e. if *S_o_* = *S_pmax_* and *S_o_* = *S_pmin_*. Such sites were determined by comparing the dominant and additive models separately for *P_max,i_* and *P_min,i_* and excluding sites found to fit the dominant model in both. It is possible that additively inherited TFBSs may be misclassified if the difference in binding intensities between the parental measurements is small enough that the F1 measurement is statistically indistinguishable from either parent due to measurement noise. To minimize this potential source of error, we restricted tested sites to those TFBSs where the difference between the means of B6_F0_ and CAST_F0_ across biological replicates was equal or greater than twice the standard deviation of the average binding intensity across biological replicates (this was set at 19 normalized counts or more). To further increase confidence in our results, we only used sites assigned to their regulatory category with BIC>1.

Over- and under-dominant TFBSs were identified by first restricting all TFBSs to those classified to a regulatory class with BIC>1. Normalized count data at each TFBS was fitted to the models described above. For each TFBS where the binding occupancy of each parent did not equalled to that of the offspring (i.e. *S_pmax_ ≠ S_o_*, *S_pmin_ ≠ S_o_*), TFBSs were classified as <u>under-dominant</u> if the mean F1 occupancy level among replicates was less than that of both parents. On the other hand, TFBSs were classified as <u>over-dominant</u> when the mean F1 occupancy level was greater than that of both parents.

### Distinguishing influences at lineage-specific TFBSs

Described below are the statistical methods used to distinguish between cis and cistrans influences at lineage-specific TFBSs. Read counts were normalized between F0 and F1 libraries as described in the previous section^73^. Lineage-specific binding sites were defined as those sites meeting these criteria: (ratio_F0_<0.05 and ratio_F1_<0.05) or (ratio_F0_>0.95 and ratio_F1_>0.95). ratio_F0_ = B6_F0_/(B6_F0_/CAST_F0_) and ratio_F1_=B6_F1_/(B6_F1_/CAST_F1_), where ratios were determined between mean levels of binding among biological replicates. We expect that a lineage-specific site that is influenced only by cis-acting variation would possess F1 count levels that are half of that in F0. Significant deviation from this 2:1 ratio would indicate variation acting in trans. We constructed the following statistical models to test the likelihood of these scenarios for each lineage-specific site and used maximum likelihood estimation and BIC to choose the model of best fit. At each TFBS, reads across replicates were modelled using the negative binomial distribution.

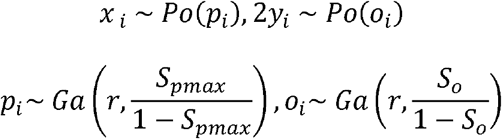

*x_i_* is defined as the normalized read count binding intensity of the variant in the *i*th F0 mouse from the strain of lineage-specific binding. *y_i_* is the binding intensity of the variant in the *i*th F1 mouse summed across both alleles. The dispersion parameter, *r*, was estimated using ‘DESeq’, as described above. We tested the two following scenarios:

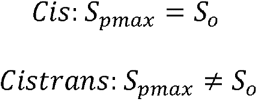

### Comparison of regulatory mechanisms underlying variation

We compared regulatory mechanisms underlying variation in gene expression, chromatin state and TF binding. Logistic regressions were used to examine the relationship between gene expression and TF binding. For each gene where expression variation was classified as affected by variation acting in cis, both cis and trans, conserved and in trans, we determined the transcriptional context by counting the numbers of each TFBSs, in each TF regulatory category, located in the window 20kb upstream and 10kb downstream of the TSS. Counts of TFBSs in each regulatory category (i.e. number of TFBSs where occupancy levels were affected by cis-acting variation, etc) were then used as four independent predictive variables. Separate regressions were performed using each of the four expression regulatory classes in turn as the dependent variable. The binary nature of the dependent variable was defined using remaining regulatory categories.

We used the same strategy to study the relationship between TF binding and chromatin state (H3k4me3), that is, the mechanistic relationship between TFBSs proximal to the histone mark was assessed using logistic regression. The size of the genomic regions used for the grouping of TFBSs was +/- 2kb from each histone mark location.

To test for shared regulatory mechanisms between H3K4me3 and gene expression, the histone marks were assigned to genes when they were located within 5 kb upstream of a TSS. Binomial tests were then used to calculate the statistical enrichment of shared regulatory mechanisms between gene expression and the associated histone marks.

We computed the diversity of TF regulatory mechanisms for genes grouped by expression mechanisms using Shannon’s diversity index (*H’*)^79^, which was calculated for each gene as follows:

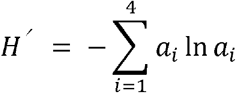

where *a_i_* is the proportion of binding sites belonging to the *i*th TF binding regulatory category within 20kb upstream or 10kb downstream of a liver-expressed protein-coding gene.

Gene expression levels show correlation with TFBS abundance, and highly expressed genes are expected to be proximal to a more diverse set of mechanisms underlying TF occupancy change than by chance alone. Hence, to control for differences in expression levels, we subsampled genes to obtain matched gene expression levels between comparison sets. Gene expression levels were compared based on the average expression value among biological replicates of the more highly expressed parent. Mean expression levels were first log transformed, then separated into 20 bins of equal consecutive intervals. Each gene affected by variation acting in both cis and trans was then matched to a conserved regulated gene assigned to the same expression bin. In the same way, genes affected by variation acting in cis were matched in expression values to conserved genes. All subsampling was done with replacement.

### Measuring inter-peak coordination of TF binding occupancy

To determine the genomic region under the influence of any set of cis-acting regulatory variants, we calculated correlation coefficients for binding intensities of TFBS pairs at successive genomic intervals away from each cis-directed TFBS. To capture the coordination of TF occupancies between TFBSs, we calculated Spearman’s correlation coefficient of allelic proportions (BL6/(BL6+CAST)) between binding sites at consecutive distance bins centred upon variants acting in cis. Spearman’s Rho was calculated for each mutually exclusive bin with their ‘anchor’ peak. Each succeeding bin was increased in interval width by one additional kb (1 kb) from the cis-acting variant. We performed linear regression using log-transformed distances as the predictor variable with Spearman’s Rho estimates as the outcome variable to quantify the decay in correlation signal (**Methods**, **Figures 4a-b, S10**).

In order for meaningful inference, we generated a null distribution of the correlation of binding strengths by comparing occupancy levels of anchor TFBSs with the occupancies of other TFBS locations sampled randomly from across the genome. Null values were calculated using TFBSs that were randomly sampled from the total pool (without replacement) to simulate a set of binned peaks for each anchor peak (anchor peaks were kept constant). The total number of binned peak simulated was equal to the total number of anchored–binned peak pairings observed. Spearman’s Rho was then calculated as described for the observed set.

To estimate the genomic distance at which the ‘elbow’ or maximum curvature of the curve occurs, we used a vector projection method on the fitted regression curve^80^. First, we drew a line connecting the points from *x* = 1kb to where *x* = 50000. Next, for every point on this line at values *x* of we extended perpendicular lines to intersect with our regression line. We then measured the lengths of each of these lines and selected the point with the longest length as the estimate of the elbow.

### Hi-C data processing and analysis

Hi-C libraries were generated from pooled liver samples from two 2-4 week old mice^49^. Raw data files were quality filtered using Trimmomatic^71^ using identical parameters to those described above. We used the Homer Hi-C software (http://homer.salk.edu/homer/interactions/) to process Hi-C reads and to identify significant interactions. Restriction sites (‘AAGCTT’) were trimmed from our reads prior to mapping to the GRCm38.p2/mm10 genome using GSNAP^74^ at a maximum of two mismatches per read. Only reads mapping to unique locations in the genome were retained. Paired reads that likely represent continuous genomic fragments or re-ligation events were removed if the reads are separated by less than 1.5x the sequencing insert fragment length (-removePEbg). Paired ends that originate from areas of unusually high read density were also removed by scanning 10kb regions in the genome and removing reads containing greater than five times the average number of reads (-removeSpikes 10000 5). Only reads where both ends of the paired read have a restriction site within the fragment length 3’ to the read were kept (-both). We also filtered reads if their ends self-ligated with adjacent restriction sites (-removeSelfLigation).

To detect significant interactions between two genomic locations, we created a background model to account for the primary sources of technical biases. For example, closely spaced loci are inevitably enriched for interactions due to their close proximity. We used Homer to normalize both for linear distance and read depth. We normalized our reads at 10 kb regions across the genome and examined the number of interactions occurring between these regions. Enrichment for significant interactions were identified using a binomial test against the expected number of interactions based on the background model that also accounts for the total number of reads mapping to each locus being tested. The parameters for the binomial test includes (i) the probability of success is the expected interaction frequency (which vary depending on restriction site locations), (ii) the number of success is the number of reads mapping between the loci, and (iii) the number of trials is the overall number of significantly interacting reads.

#### Data availability

Raw data have been deposited under ArrayExpress accession E-MTAB-4089. Processed data are available from http://www.ebi.ac.uk/research/flicek/publications/FOG19.

## ACKNOWLEDGEMENTS

We thank the CRUK - CI Genomics, BRU, and Bioinformatics Cores for technical assistance and the EMBL-EBI systems team for management of computational resources. This research was supported by the European Molecular Biology Laboratory (E.S.W., D.T., J.M., P.F.), Cancer Research UK (B.M.S., T.R., F.C., C.F., A.R., D.T.O.), the BOLD ITN (B.M.S.), Darwin Fellowship (AK), the Wellcome Trust (WT202878/B/16/Z, WT108749/Z/15/Z) (P.F.), (WT202878/A/16/Z) (D.T.O), (WT095606) (AFS) and (WT098051) (P.F., D.T.O.), EMBO Long-term (ALTF1518-2012) and Advanced Fellowships (aALTF1672-2014) (E.S.W.), and by the European Research Council (award 615584) and EMBO Young Investigator Programme (D.T.O.).

## AUTHOR CONTRIBUTIONS

ESW, BMS, DT, AFS, JCM, DTO, PF designed experiments; BMS, AR, AK performed wet lab experiments; ESW performed computational analyses; BMS, ESW, AK, TR collated and quality controlled the data; FC, CF provided experimental assistance; ESW, BMS, PF, DTO wrote the manuscript; DTO, PF oversaw the work.

